# Antigenic mapping of the hemagglutinin of the H9 subtype influenza A viruses using sera from Japanese quail (*Coturnix c. japonica*)

**DOI:** 10.1101/2023.05.18.541344

**Authors:** Silvia Carnaccini, C. Joaquín Cáceres, L. Claire Gay, Lucas M. Ferreri, Eugene Skepner, David F. Burke, Ian H. Brown, Ginger Geiger, Adebimpe Obadan, Daniela S. Rajao, Nicola S. Lewis, Daniel R. Perez

## Abstract

Influenza A viruses (FLUAV) of the H9N2 subtype are zoonotic pathogens that cause significant economic damage to the poultry industry. Vaccination to prevent and control H9N2 infections in poultry is widely employed in the Middle East and Asia. We used phylogenetics and antigenic analysis to study the antigenic properties of the H9 hemagglutinin (HA) using sera produced in Japanese quail (*Coturnix c. japonica*). Consensus HA1 sequences were generated to capture antigenic diversity among isolates. We constructed chimeric H9N2 viruses containing the HA1 of each consensus sequence on a constant isogenic backbone. The resulting viruses were used to generate antisera from quail, a common and significant minor poultry species whose anti-HA response profiles remain poorly defined. Antigenic maps were generated by plotting the cross-hemagglutination inhibition (HI) data from the panel of quail sera against the chimeric constructs and 51 H9 field isolates. The chimeric antigens were divided into four different antigenic profiles (cyan, blue, orange, and red). Site-directed mutagenesis analysis showed 9 amino acid positions of antigenic relevance. Substitutions at amino acid positions 149, 150, and 180 (H9 HA numbering) had relatively significant impact on HI activity using quail sera. Substitutions E180A and R131K/E180A led to the most significant antigenic change transitions. This study provides insights into the antigenic profile of H9 FLUAVs, with important implications for understanding antigenic drift and improving vaccine development for use in minor poultry species.

**IMPORTANCE:** Determining the relevant amino acids involved in antigenic drift on the surface protein hemagglutinin (HA) is critical to understand influenza virus evolution and efficient assessment of vaccine strains relative to current circulating strains. We used antigenic cartography to generate an antigenic map of the H9 HA using sera produced in one of the most relevant minor poultry species, Japanese quail. Key antigenic positions were identified and tested to confirm their impact on the antigenic profile. This work provides a better understanding of the antigenic diversity of the H9 HA as it relates to reactivity to quail sera and will facilitate a rational approach for selecting more efficacious vaccines against poultry-origin H9 influenza viruses in minor poultry species.

## INTRODUCTION

Influenza A virus (FLUAV) of the H9N2 subtype are enzootic in poultry in Asia, the Middle East, and parts of Africa, where they cause significant economic losses to the poultry industry due to high morbidity and mortality of poultry flocks (1, 2). More importantly, Eurasian-origin H9N2 FLUAVs are zoonotic viruses and they have provided the internal gene constellation to more virulent zoonotic strains, notably the H5N1 Guangdong lineage, the Asian-lineage H7N9, H10N8 and H3N8 FLUAVs (3-5). The World Health Organization (WHO) has placed H9N2 FLUAVs among those with pandemic concern. Recently, we proposed a consistent numerical nomenclature for the HA of the H9 subtype, similar to the system adopted for the H5 subtype (2). Initially, two major geographically distinct H9 lineages were identified: the American (h9.1) and the Eurasian (h9.2) lineages (2). The continuous circulation of H9 FLUAVs in poultry in Asia has led to significant evolution and, consequently, phylogeographic diversity among the Eurasian lineage viruses leading to several sub-lineages and sub-sub-lineages. Currently, Eurasian H9 HA sequences fall into three major sub-lineages: h9.2 (previously referred to as Y439, prototype dk/HK/Y439/1997), h9.3 (BJ94, prototype ck/Bei/1/94), and h9.4 (G1, prototype qa/HK/G1/1997). The H9.2 sub-lineage can be divided further into several sub-sub-lineages, including h9.2.1 and h9.2.2, which are mostly found in wild birds, and h9.2.3, also known as Korean-strict and found in poultry in South Korea. The h9.3 sub-lineage is particularly prominent in China and Southeast Asia, with the presence of at least 9 sub-sub-lineages, h9.3.1-h9.3.9. Previous studies suggested dividing the h9.4 sub-lineage into Eastern and Western sub-sub-lineages based on their respective geographic prevalence (2). However, due to early indications of incongruent geographic boundaries among the Eastern and Western h9.4 strains, we proposed an alternative numerical nomenclature, h9.4.1 and h9.4.2, respectively (3, 4).

To prevent and control H9N2 virus infections in poultry, several countries in Asia and Middle East have resorted to vaccination programs (5-11). Antigenic drift of H9 FLUAVs is readily observed in the field, likely a combination of natural evolution and vaccine use (5-11). Near and around the receptor binding site, the globular head HA1 portion of the H9 HA contains two partially overlapping antigenic sites. These sites have been defined previously using mouse monoclonal antibodies (mAbs) and are known as sites I and II or, more recently, as sites H9-A and H9-B, respectively (12-16). Site H9-A is immunodominant compared to site H9-B (13, 17). A limited set of the most prominent poultry-adapted Eurasian lineages from specific regions have been examined antigenically (12-14, 18-20). Most antigenic analyses of H9N2 viruses have been performed using chicken sera and, to a lesser extent, ferret sera, but not with sera from minor poultry species such as quail. Japanese quails have been suggested as key players in the genesis of influenza viruses with respiratory tract tropism (21, 22). Quail show wide distribution in the respiratory tract of both avian-like (SAα2.3) and human-like (SAα2.6) sialic acid receptors, which may have contributed to the emergence of the poultry adapted H9N2 strains with human-like receptor preference (23, 24). Anti-H9 sera have been raised by different approaches and regimes, which act as confounding factors to assess antigenicity faithfully (17, 25-28). Immunization approaches have included either live virus challenge or most typically inactivated/adjuvanted viruses in either single or prime and boost infection or vaccination. Despite the absence of a standardized approach for sera production, these analyses have shown some significant clues about the antigenic makeup of the H9 HA. Combined with studies using mouse mAbs, a cluster of amino acids has been shown to affect the antigenic profile of the HA, namely those at positions 72, 74, 121, 131, 135, 150, 180, 183, 195, 198, 216, 217, 249, 264, 276, 288, and 306 (H9 numbering throughout the manuscript)(17, 26, 27, 29, 30). Further analyses on the contributions of each of these and alternative positions to antigenicity/receptor binding avidity are discussed later in the context of this report’s findings.

To broaden the understanding of the antigenic diversity of HAs of H9 FLUAVs, we included strains from the American and Eurasian lineages. Starting from an initial phylogenetic analysis of nucleotide sequences corresponding to the HA1 region of the HA, we identified 18 clades utilizing sequence information of strains from 1966 to 2020. Analyses of these clades led to the selection of 10 consensus sequences that largely embodied the amino acid diversity within each H9 lineage/sub-lineage/sub-sub-lineage. The 10 HA1 sequences were used to generate chimeric H9 HA gene segments carrying a constant HA2 portion derived from the prototypic strain gf/HK/WF10/1999 (H9N2) (WF10) (31, 32). The chimeric HA constructs were subsequently used for reverse genetics. To better understand the H9 HA antigenic make-up in the context of neutralizing responses in minor poultry, Japanese quails were challenged with the chimeric H9 HA viruses. Anti-H9 quail sera were used to perform hemagglutination inhibition (HI) assay and antigenic cartography (15, 33). These analyses showed H9 HA antigens positioned in 4 antigenic clusters in the antigenic map, with additional outliers. Viruses carrying amino acid substitutions at relevant antigenic positions were generated to explain cluster transitions. These results provide new insights into the antigenic evolution of H9N2 influenza viruses and offer new opportunities to improve vaccine development.

## RESULTS

### Phylogenetic analysis, consensus sequences, and antigenically relevant amino acids on H9 HA

Using the H9 HA1 region a maximum likelihood phylogenetic tree was established based on nucleotide sequences from isolates between 1966 to 2016 and then updated with sequences up to 2020. The phylogenetic analysis allowed the identification of different clades (h9.1.1 to h9.4.2). Consensus sequences were generated for each clade, n=10 **(Fig 1A)**. The % amino acid identity ranged from 83.1% (h9.2.3 vs. h9.3.9) to 98.4% (h9.3.3 vs. h9.3.4). The number of amino acid differences in the HA1 region between the consensus sequences and the HA of the prototypic h9.4.1 strain WF10 were 31 (h9.4.2), 35 (h9.2.4), 36 (h9.3.3), 37 (h9.2.2), 38 (h9.3.4), 39 (h9.1.1 and h9.3.3), 44 (h9.3.7), 47 (h9.2.3), and 48 (h9.3.9), respectively **(Fig 1B)**. Chimeric HA constructs were used for reverse genetics in the WF10 backbone. In addition to the wild-type WF10 strain, 8 out of the 10 chimeric HA constructs resulted in viable H9N2 viruses. No virus rescue was obtained for the chimeric HA representing the h9.2.3 and h9.2.4 clades. Analysis of the HA1 portion of the consensus viruses and the closest relative from a subset of field viruses showed high similarity **(Fig 1B)**. For WF10, the closest relative was A/qa/HK/G1/97 (98.4%); for h9.4.2, A/ck/Pak/47/03 (98.9%); for h9.3.9 and h9.3.7, A/dk/Hunan/1/2006 (93.3% and 96.5%, respectively); for h9.3.4, A/dk/HK/Y280/97 (96.9%); and for h9.3.3, A/ck/Sichuan/5/97 (98.5%). The % of identity between h9.2.2 and A/ml/Fin/Li13384/2010 and h9.1.1 and A/rt/New Jersey/AI11-1946/2011 was 98.6% and 95.5% respectively.

**Fig. 1.**
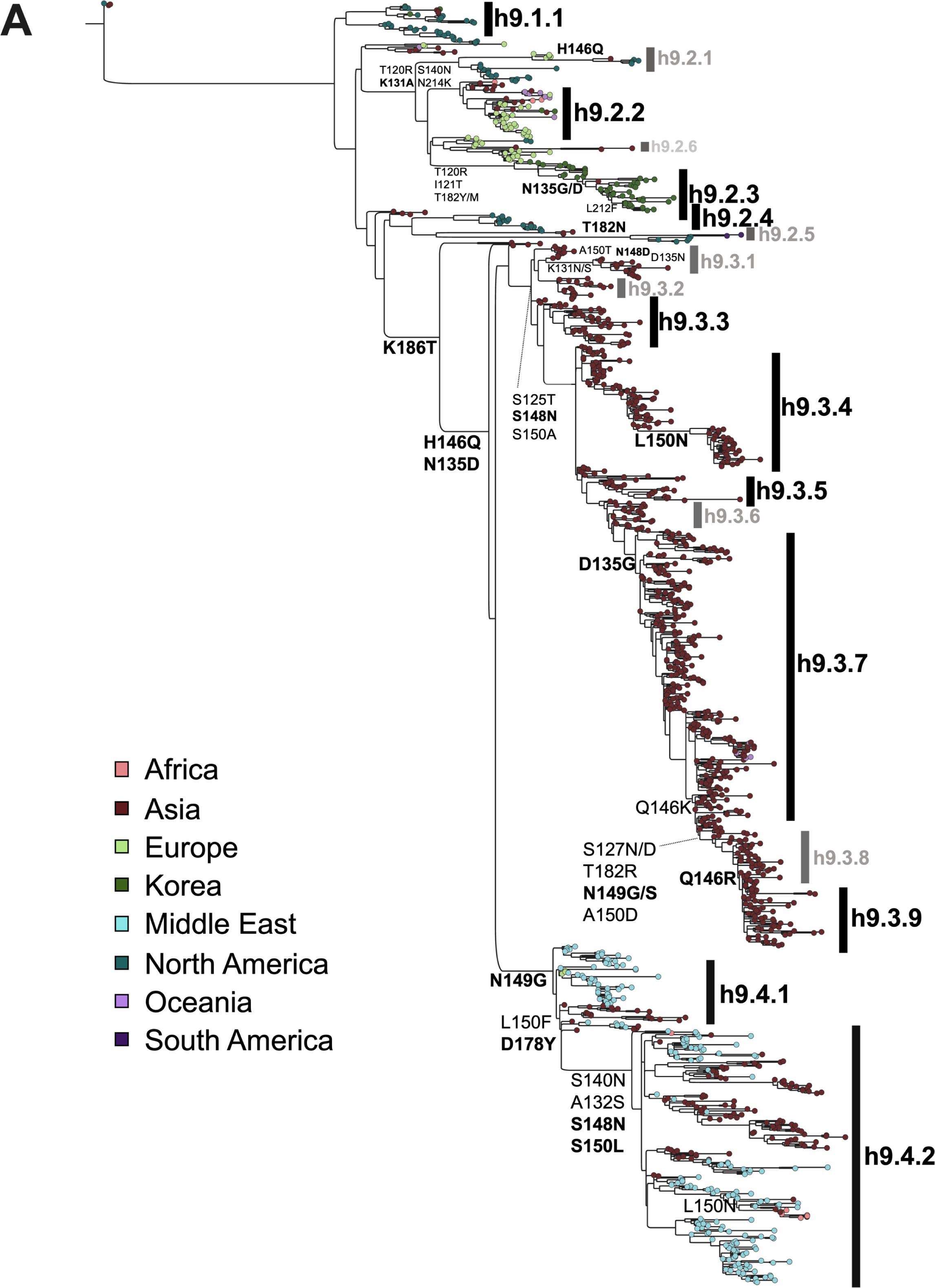
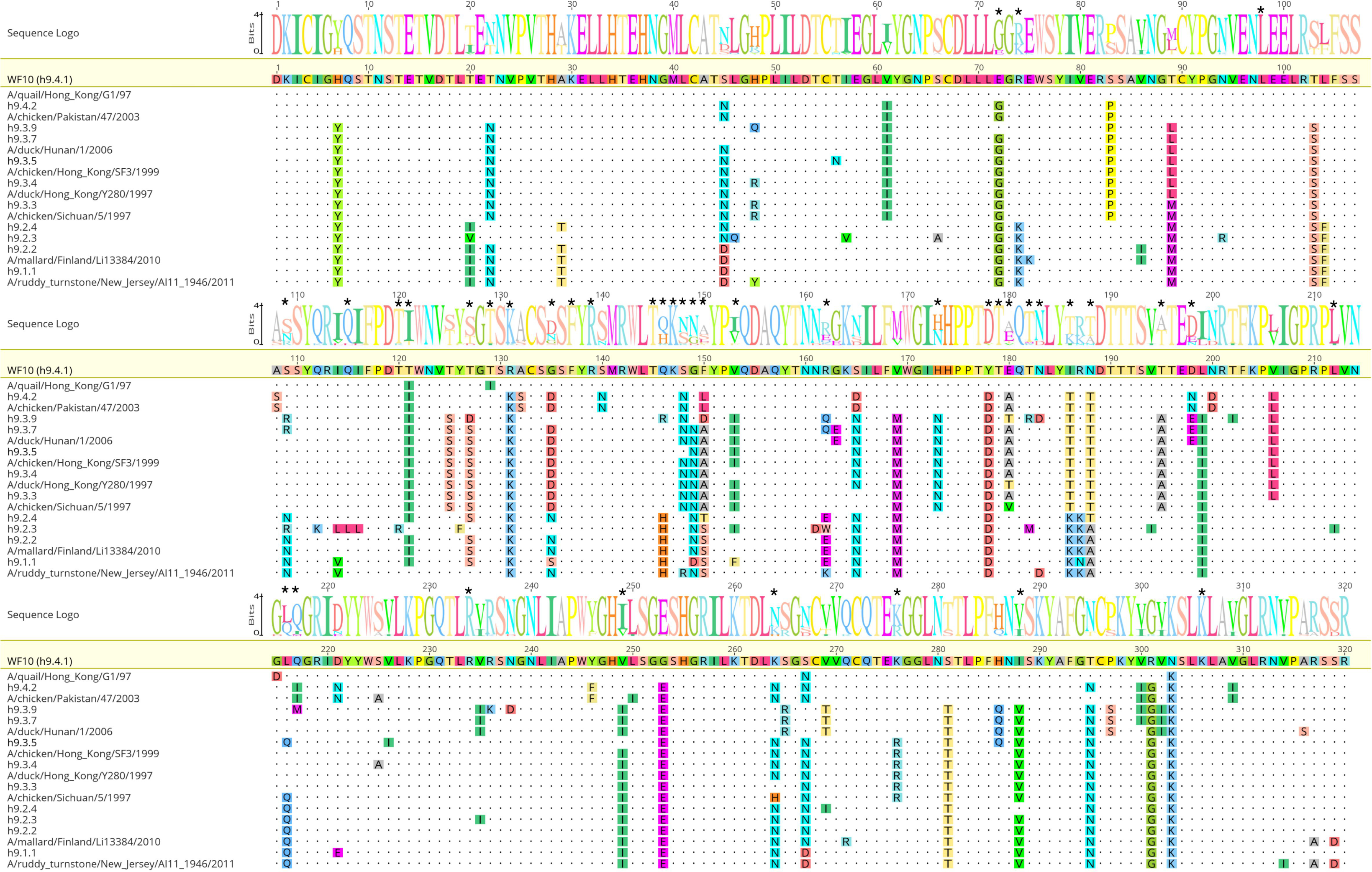
Global phylogenetic analysis of H9N2 FLUAV. **(A)** Maximum Likelihood phylogeny of 1,316 H9 avian HA1 nucleotide sequences from the GISAID and IRD databases updated July 14^th^, 2020, generated with RaxML followed by Garli branch length optimization. Nodes at the end of each branch are color-coded based on the geographic origin of each isolate. Amino acid substitutions using one-letter code and numbering based on H9 HA mature sequence are shown. Highlighted in black are H9 sub-lineages chosen to generate consensus HA1 region sequences and to produce chimeric H9 HA constructs with a constant HA2 region. Sub-lineages that were unsuccessful in reverse genetics are shown in grey. The h9.4.1 consensus is represented by the prototypic virus A/gf/HK/WF10/1999 (H9N2). **(B)** WebLogo by Geneious v2022.2.2 with the alignment of the consensus HA1 amino acid sequences and closest relatives in each case (under each consensus sequence) against WF10 wild-type HA1. (*) on top of amino acid positions indicate potentially relevant antigenic amino acids. The closest relative for h9.3.9 and h9.3.7 is the same (A/dk/Hunan/1/2006). No closest relative against h9.2.4 and h9.2.3 are shown since no viable virus was obtained for those clades.

### HI responses against consensus clades viruses in quail

To generate antisera against the chimeric HA consensus viruses, we chose Japanese quail (*Coturnix c. japonica*) as a relevant minor poultry host of H9 FLUAVs (21, 31). Groups of quail (9 groups, n=6/group) were inoculated with either of each H9N2 chimeric virus or WF10 wild type **(Fig 2A).** At 14 days post-inoculation (14 dpi), quail were boosted subcutaneously with inactivated-adjuvanted preparations of each virus. At 28 dpi, quail were terminally bled, and 2 independent pooled sera were generated (3 birds per pool). We analyzed the seroconversion to the homologous virus in inoculated quail by HI assays showing titers between 1280 and 5120 against the homologous viruses **(Table 1)**. The highest homologous HI titers were obtained for h9.3.3 and h9.3.9, with a titer of 5120 in each case. Similarly, titers of 2560-5120 were observed for h9.3.4, while a titer of 2560 was obtained for h9.4.2. In the case of h9.3.7 and WF10, titers of 1280-2560 were observed. The h9.1.1 and h9.2.2 groups were the exception, with HI titers of 80-160 and 40-160, respectively, which are considerably lower than the other consensus viruses. Taken together, the homologous HI data shows high levels of neutralizing antibodies against the different consensus viruses, except h9.1.1 and h9.2.2, which elicit poor antibody responses in the quail model.

**Fig. 2.**
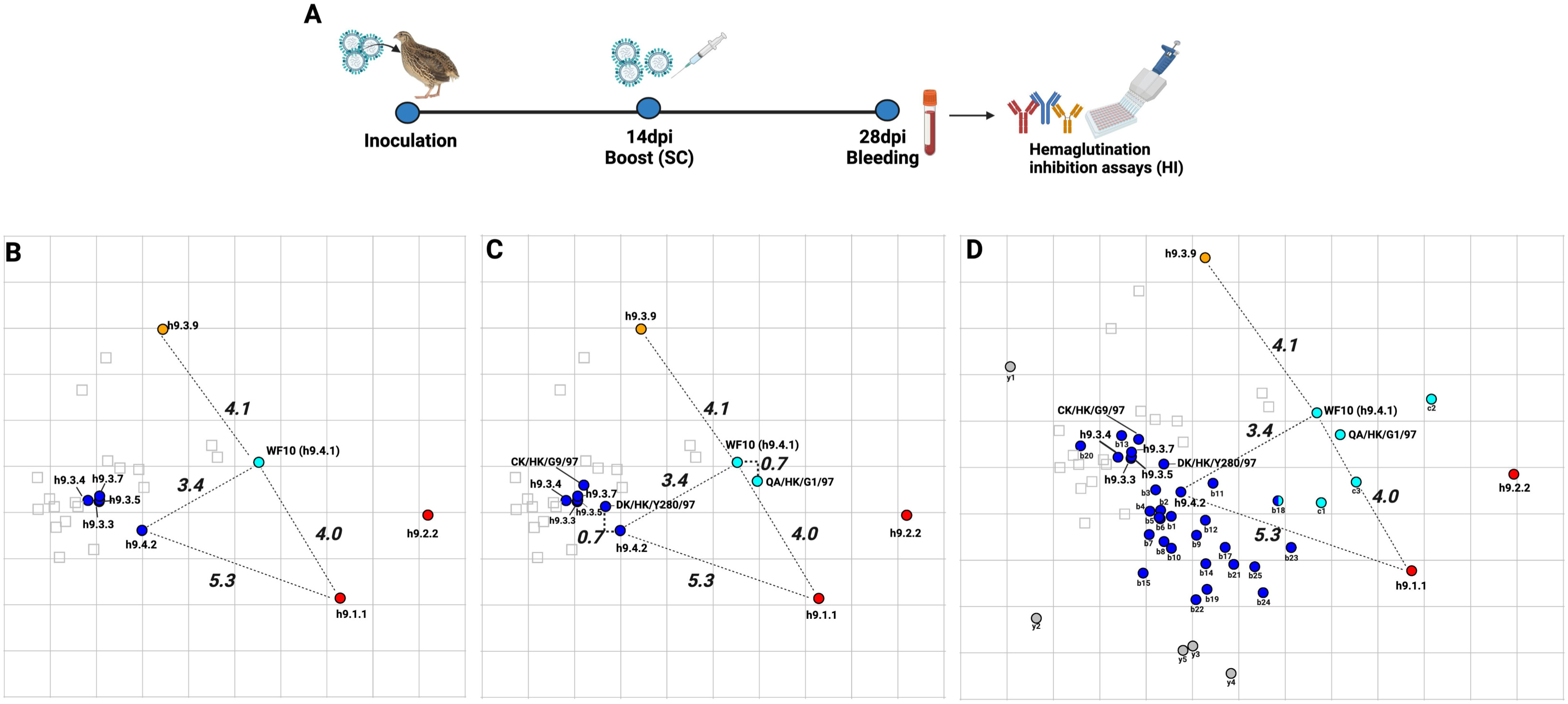
Antigenic maps using quail sera against H9 viruses. **(A)** Schematic representation of the inoculation in quails produced with Biorender.com. Birds were inoculated with each consensus virus at day 0, and at 14dpi they were boosted with homologous inactivated-adjuvanted virus preparations. At 28dpi, quails were bled, and the antisera were obtained. **(B)** Antigenic map with spheres representing consensus viruses and squares representing the different antisera. Viruses are highlighted and colored by respective clusters (cyan, red, blue, and orange). AU distances between representative antigens from each cluster are shown next to dashed grey lines connecting them. **(C)** Antigenic map with spheres representing consensus viruses + prototypical strains (QA/HK/G1/07, DK/HK/Y280/97, and CK/HK/G9/97) and squares representing the different antisera. Viruses are highlighted and colored by respective clusters (cyan, red, blue, and orange). AU distances between representative antigens from each cluster and prototypic strains are shown next to dashed grey lines connecting them. **(D)** Antigenic map with spheres representing consensus viruses + field isolates (n=46) and squares representing the different antisera. Viruses are highlighted and colored by respective clusters (cyan, red, blue, and orange). AU distances between representative antigens from each cluster are shown next to dashed grey lines connecting them. Except for the orange antigenic h9.3.9 antigen, all other antigens that showed sera reactivity but did not fall into an antigenic cluster are shown in grey as outliers. Specific viruses are denoted by codes shown in Table 2.

**Table 1.**
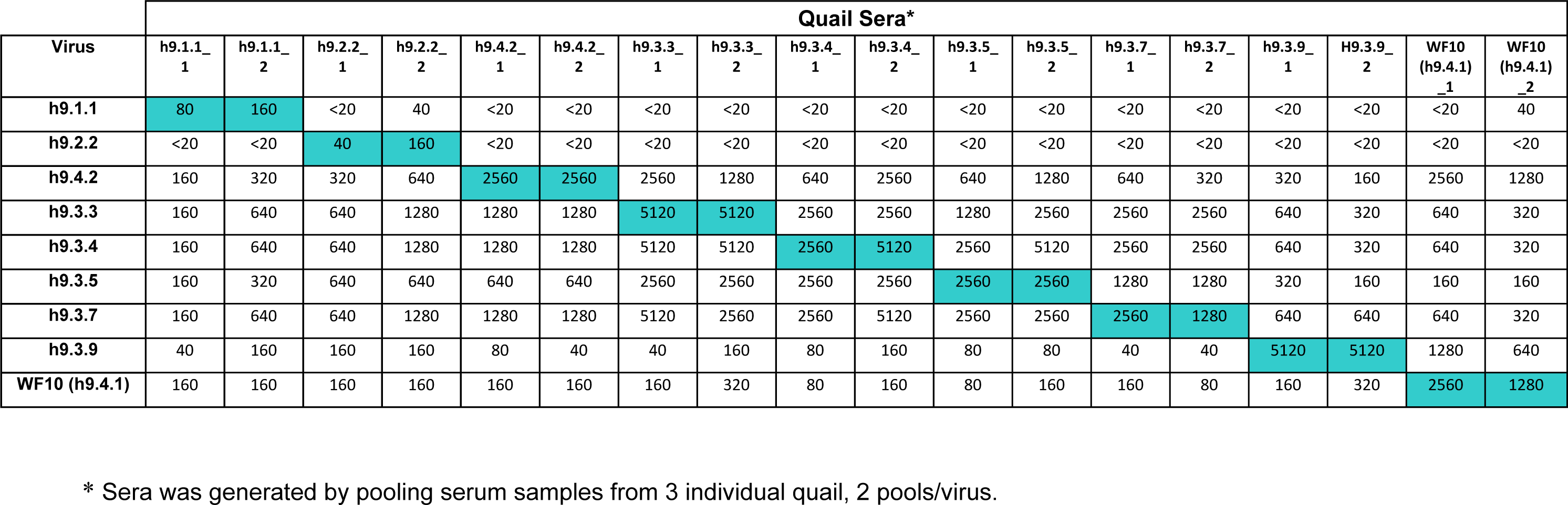
Cross-HI titers against chimeric HA-WF10 (H9N2) viruses using quail sera.

**Table 2.** Cluster location and antigenic distances of consensus viruses and field isolates.

| **Virus** | **Closest Centroid** | **Antigenic distance** | **Label** |
| --- | --- | --- | --- |
| **h9.1.1** | **h9.1.1** | **0** | **h9.1.1** |
| **h9.2.2** | **h9.1.1** | 2.9 | **h9.2.2** |
| A/ma/Finland/LI13384/2010c | **h9.1.1** | 2.6 | **r1** |
| A/rt/New Jersey/AI11-1946/2011 | **h9.1.1** | 0.9 | **r2** |
| A/sh/Delaware/9/1996 | **h9.1.1** | 2.0 | **r3** |
| A/ph/Republic of Ireland/PV18/1997 | **h9.1.1** | 0.3 | **r4** |
| A/dk/Hong Kong/702/1979 | **h9.1.1** | 1.4 | **r5** |
| A/sh/Delaware/277/1999 | **h9.1.1** | 0.5 | **r6** |
| A/tk/USA/6707-1/1996 | **h9.1.1** | 1.2 | **r7** |
| A/ck/Shijiazhuang/2/1999 | **h9.1.1** | 1.0 | **r8** |
| A/qa/Arkansas/29209/1993 | **h9.1.1** | 1.3 | **r9** |
| A/ck/Sichuan/5/1997 | **h9.1.1** | 2.7 | **r10** |
| A/ck/Jordan/554/2003 | **h9.1.1** | 1.7 | **r11** |
| **h9.4.2** | **h9.4.2** | **0** | **h9.4.2** |
| **h9.3.3** | **h9.4.2** | 1.3 | **h9.3.3** |
| **h9.3.4** | **h9.4.2** | 1.5 | **h9.3.4** |
| **h9.3.5** | **h9.4.2** | 1.3 | **h9.3.5** |
| **h9.3.7** | **h9.4.2** | 1.4 | **h9.3.7** |
| A/ck/Hong Kong/G9/1997 | **h9.4.2** | 1.4 | **G9** |
| A/dk/Hong Kong/Y280/1997 | **h9.4.2** | 0.7 | **Y280** |
| A/ck/Bangladesh/301/2007 | **h9.4.2** | 0.6 | **b1** |
| A/ck/Saudi Arabia/3489V08-50/2008 | **h9.4.2** | 0.6 | **b2** |
| A/ck/Tunisia/812/2012 | **h9.4.2** | 0.5 | **b3** |
| A/ck/Pakistan/47/2003 | **h9.4.2** | 0.8 | **b4** |
| A/ck/Saudi Arabia/3489V08-44/2006 | **h9.4.2** | 0.7 | **b5** |
| A/ck/Nepal/PT22/2013 | **h9.4.2** | 0.7 | **b6** |
| A/ck/Bangladesh/262-O-SUN-1/2016 | **h9.4.2** | 1.1 | **b7** |
| A/ck/Saudi Arabia/A/2010 | **h9.4.2** | 1.1 | **b8** |
| A/ck/Tunisia/12/2010 | **h9.4.2** | 1.0 | **b9** |
| A/ck/India/1/2003 | **h9.4.2** | 1.2 | **b10** |
| A/ck/Germany/K1009/1998 | **h9.4.2** | 0.7 | **b11** |
| A/ck/Pakistan/UDL 7/2008 | **h9.4.2** | 0.8 | **b12** |
| A/ck/Hong Kong/SF3/1999 | **h9.4.2** | 1.7 | **b13** |
| A/ck/Iran/AIV 1/2003 | **h9.4.2** | 1.6 | **b14** |
| A/ck/Iraq/30/2011 | **h9.4.2** | 1.9 | **b15** |
| A/ck/Afghanistan/329V09/2008 | **h9.4.2** | 1.5 | **b17** |
| A/qa/United Arab Emirates/302/2001 | **h9.4.2** | 2.1 | **b18** |
| A/ck/Saudi Arabia/3489V08/2005 | **h9.4.2** | 2.2 | **b19** |
| A/dk/Hunan/1/2006 | **h9.4.2** | 2.4 | **b20** |
| A/av/Middle East/2/1998 | **h9.4.2** | 1.9 | **b21** |
| A/ck/Saudi Arabia/3489V08-47/2007 | **h9.4.2** | 2.3 | **b22** |
| A/ck/Nepal/1-220/2013 | **h9.4.2** | 2.7 | **b23** |
| A/ck/Guangdong/11/1997 | **h9.4.2** | 2.8 | **b24** |
| A/ty/Israel/1266/2003 | **h9.4.2** | 2.3 | **b25** |
| **A/guinea fowl/Hong Kong/WF10/1999** | **WF10** | **0** | **WF10** |
| A/qa/Hong Kong/G1/1997 | **WF10** | **1** | **G1** |
| A/ck/Saudi Arabia/S11A/2003 | **WF10** | 2.4 | **c1** |
| A/ty/Netherlands/11015452/2012 | **WF10** | - | **c2** |
| A/qa/Hong Kong/A28945/88 | **WF10** | 1.7 | **c3** |
| **h9.3.9** | **h9.3.9** | **-** | **orange** |
| A/ck/Beijing/8/1998 | - | - | **y1** |
| A/ck/Hebei/3/1998 | - | - | **y2** |
| A/ck/United Arab Emirates/H4TR/2011 | - | - | **y3** |
| A/ck/Libya/D31 TRACH/2006 |  |  | **y4** |
| A/ck/Jordan/901-F5/2003 |  |  | **y5** |
| **A/ck/North_Korea/99029/1999** | - | - | **Black** |
| **A/ck/Tunisia/345/2011** | - | - | **Black** |
| **A/dk/Hong_Kong/448/1978** | - | - | **Black** |
| **A/mallard/England/7798_6499/2006** | - | - | **Black** |
| **A/ql/Saudi_Arabia/489_46v08/2006** | - | - | **Black** |

**a**Determined by antigenic analysis through ACMACS. **b**One unit of antigenic distance is equal to a 2-fold difference in the HI assay
**c**All virus strains are of the H9N2 subtype except where noted. \*Not Tested; 1\*Non cross reactive against the panel of antisera Animal species acronyms dl=duck. ck=chicken, ty=turkey, ph=pheasant, qa=quail, mal=mallard, rt= ruddy turnstone, sh=shorebird, av=avian.

### Antigenic analysis of H9 HA

Using the antigenic cartography platform, the cross-HI data obtained were merged and visualized by generating maps in which the spheres represent antigens and the squares the sera, distributed into space. Antigenic distances between antigens in the map are expressed in antigenic units (AU, 1AU corresponds to a 2-fold dilution of antiserum in the HI assay). Dimensional analysis of the HI dataset led to lower error yield in the 3D maps, though 2D maps were selected for better visualization, given that the relationship between consensus antigens remained unvaried. The antigens were grouped into 4 different clusters as described in Material and Methods **(Fig 2B)**. We used 3 AU or a ≥8-fold loss in cross-reactivity, as defined for the human seasonal vaccine strain update (WHO recommendation), as the threshold of significant antigenic difference. The WT WF10 HA prototypic h9.4.1 antigen (cyan) was 3.4 AU from the h9.4.2 antigen (blue). The h9.3.3, h9.3.4, h9.3.5, and h9.3.7 antigens (blue) clustered antigenically very close to each other (<0.3 AU) and with 1.3, 1.6, 1.3 and 1.4 AU from the h9.4.2 blue antigen respectively. The h9.3.9 antigen (orange) was 4.5 AU from the h9.3.7 consensus (blue), the closest phylogenetic relative, and 5.1 AU from the h9.4.2 blue antigen. The distance between WT WF10 HA prototypic h9.4.1 antigen (cyan) and the h9.3.9 antigen (orange) was 4.1. The h9.1.1 and h9.2.2 consensus antigens (red) showed relatively close antigenic relationships (2.9 AU from each other), but distances between h9.1.1 and WF10 (cyan), h9.4.2 (blue), and h9.3.9 (orange) antigens were 4.0, 5.3, and 8.1, respectively. It must be noted that the robustness of positioning of h9.1.1 and h9.2.2 must be interpreted cautiously due to the relatively low inherent antigenicity/immunogenicity compared to the rest of the consensus antigens.

To better define whether the consensus chimeric H9 HA viruses captured the antigenic profile of prototypic strains within each clade, the quail sera was used in HI assays using a subset of closest prototypical field strains available **(Fig 2C and table 2)**. The positioning of the prototypic field antigens relative to the consensus antigens was generally consistent with their position in the phylogenetic tree. The prototypic A/qa/HK/G1/97 (h9.4.1) antigen was 0.7 AU from the WF10 h9.4.1 antigen (cyan). Two prototypic strains, A/ck/HK/G9/1997 (G9, h9.3.3-like) and A/dk/HK/Y280/1997 (Y280, h9.3.4-like), clustered together with the h9.3.3, h9.3.4, h9.3.5, h9.3.7, and h9.4.2 consensus sequences as part of a blue cluster. The antigenic distances between G9 and h9.3.3 were 0.4 AU and 1.5 AU between G9 and h9.4.2, suggesting that genetically similar viruses are also antigenically similar. In the case of Y280, 1.0 AU of difference was observed from h9.3.4 and 0.7 AU from h9.4.2.

We expanded these analyses to 48 additional field strains **(Fig 2D),** bringing the panel to 51 field strains **(Table S1)**. The analysis of the other consensus viruses and the antigenic distances from their closest relatives **(Fig 1B)** revealed similarities between genetic and antigenic properties except for h9.3.3 and h9.3.9 due to the distances between the consensus viruses and their respective closest relatives **(Table 2).** Distances between h9.3.3 and A/ck/Sichuan/5/1997 were 5.7 AU, while distances between h9.3.9 and A/dk/Hunan/1/2066 were 4.9 AU placing consensus viruses and closest relatives in different clusters. The remaining consensus showed a good correlation with their closest relatives with distances between h9.1.1-A/rt/New Jersey/AI11-1946/2011, h9.2.2-A/ma/Li13384/2010, h9.3.5-A/ck/HK/SF3/99, h9.3.7-A/dk/Hunan/1/2006 and h9.4.2-A/ck/Pakistan/47/2003 of 0.9 AU, 2.4 AU, 0.5 AU, 1.1 AU, and 0.8 AU respectively. From the 51 field isolates evaluated **(Fig 2D)**, 11 fell within the red cluster, 26 within the blue cluster, 4 within the cyan cluster, and none in the orange cluster **(Table 2)**. Due to the low reactivity of the antigenicity/immunogenicity of the red-cluster consensus viruses (h9.1.1 and h9.2.2) compared to the rest of the consensus antigens, field isolates of the red cluster were removed from the map **(Fig 2D)**. The h9.3.9 antigen was antigenically distinct from the rest of the h9.3 lineage viruses with AU distances of 4.1 (h9.3.3), 4.3 (h9.3.4), 4.1 (h9.3.5), and (h9.3.7). Further, none of the field isolates evaluated fell within 3 AU of distance from h9.3.9. The closest antigen to h9.3.9 was WF10 (cyan, 4.1 AU) **(Table 3).** A/ck/Beijing/8/1998 (h9.3.3), A/ck/Hebei/3/1998 (h9.3.3), A/ck/UAE/H4TR/2011 (h9.2.2), A/ck/Libya/D31 TRACH/2006, A/ck/Jordan/901-F5/2003 (h9.4.1, G1-like) were classified as outliers as they were >3.0 in AU distance from any of the consensus antigens (grey; **Fig. 2 and Table 2**). H9s with <40 HI titers against any of the antisera were considered to have low to no cross-reactivity against any of the antisera and were removed from the antigenic analysis **(Table 2)**. Overall, we observed mismatching between phylogenetic and antigenic analysis among viruses within the h9.3 and h9.4 lineages, mostly poultry isolates. Both h9.3 and h9.4 phylogenetic lineages contained the most antigenically variable strains, which fell under the different clusters (and some were outliers). The A/qa/UAE/302/2001 (b18, **Fig 2D**) HA antigen was equally distant from h9.4.2 and WF10 antigens with 2.1 AU of distance in both cases **(Table 2)**. Taken together, the results provide an antigenic map of the H9 HA using consensus and wild type HA sequences probed with quail sera.

**Table 3.** Antigenic distances between consensus viruses and different field isolates.

|  | WF10 | h9.4.2 | h9.3.3 | h9.3.4 | h9.3.5 | h9.3.7 | h9.3.9 | h9.1.1 | h9.2.2 |
| --- | --- | --- | --- | --- | --- | --- | --- | --- | --- |
| **A/guinea fowl/Hong Kong/WF10/1999** | 0.0 | 3.4 | 4.1 | 4.3 | 4.1 | 4.1 | 4.1 | 4.0 | 4.5 |
| h9.4.2 | 3.4 | 0.0 | 1.3 | 1.6 | 1.3 | 1.4 | 5.1 | 5.3 | 7.2 |
| h9.3.3 | 4.1 | 1.3 | 0.0 | 0.3 | 0.1 | 0.1 | 4.6 | 6.5 | 8.3 |
| h9.3.4 | 4.3 | 1.6 | 0.3 | 0.0 | 0.3 | 0.3 | 4.7 | 6.8 | 8.5 |
| h9.3.5 | 4.1 | 1.3 | 0.1 | 0.3 | 0.0 | 0.1 | 4.6 | 6.5 | 8.3 |
| h9.3.7 | 4.1 | 1.4 | 0.1 | 0.3 | 0.1 | 0.0 | 4.5 | 6.6 | 8.3 |
| h9.3.9 | 4.1 | 5.1 | 4.6 | 4.7 | 4.6 | 4.5 | 0.0 | 8.1 | 8.1 |
| h9.1.1 | 4.0 | 5.3 | 6.5 | 6.8 | 6.5 | 6.6 | 8.1 | 0.0 | 2.9 |
| h9.2.2 | 4.5 | 7.2 | 8.3 | 8.5 | 8.3 | 8.3 | 8.1 | 2.9 | 0.0 |
| **A/av/Middle East/2/1998** | 3.7 | 1.9 | 3.2 | 3.4 | 3.2 | 3.3 | 6.6 | 3.8 | 6.3 |
| **A/ck/Afghanistan/329V09/2008** | 3.5 | 1.5 | 2.8 | 3.0 | 2.8 | 2.9 | 6.3 | 4.1 | 6.4 |
| **A/ck/Bangladesh/262-O-SUN-1/2016** | 4.5 | 1.1 | 1.7 | 1.8 | 1.7 | 1.8 | 6.1 | 5.7 | 8.0 |
| **A/ck/Bangladesh/301/2007** | 3.9 | 0.6 | 1.5 | 1.7 | 1.5 | 1.6 | 5.6 | 5.3 | 7.4 |
| **A/ck/Beijing/8/1998** | 6.7 | 4.6 | 3.3 | 3.0 | 3.3 | 3.2 | 4.8 | 9.7 | 11.1 |
| **A/ck/Germany/K1009/1998** | 2.7 | 0.7 | 1.8 | 2.1 | 1.8 | 1.9 | 4.9 | 4.7 | 6.5 |
| **A/ck/Guangdong/11/1997** | 4.1 | 2.8 | 4.1 | 4.3 | 4.1 | 4.2 | 7.3 | 3.2 | 6.0 |
| **A/ck/Hebei/3/1998** | 7.5 | 4.1 | 4.0 | 3.9 | 4.0 | 4.1 | 8.6 | 8.2 | 10.8 |
| **A/ck/Hong Kong/G9/1997** | 3.9 | 1.5 | 0.4 | 0.6 | 0.4 | 0.3 | 4.2 | 6.5 | 8.1 |
| **A/ck/Hong Kong/SF3/1999** | 4.2 | 1.8 | 0.5 | 0.5 | 0.5 | 0.4 | 4.2 | 6.9 | 8.5 |
| **A/ck/India/1/2003** | 4.3 | 1.2 | 2.1 | 2.3 | 2.2 | 2.2 | 6.3 | 5.2 | 7.6 |
| **A/ck/Iran/AIV 1/2003** | 4.0 | 1.6 | 2.8 | 3.0 | 2.8 | 2.9 | 6.6 | 4.4 | 6.9 |
| **A/ck/Iraq/30/2011** | 5.1 | 1.9 | 2.5 | 2.6 | 2.5 | 2.6 | 6.9 | 5.8 | 8.2 |
| **A/ck/Jordan/554/2003** | 3.2 | 3.5 | 4.8 | 5.1 | 4.8 | 4.9 | 7.1 | 1.8 | 4.4 |
| **A/ck/Jordan/901-F5/2003** | 5.9 | 3.4 | 4.3 | 4.4 | 4.3 | 4.4 | 8.5 | 5.2 | 8.1 |
| **A/ck/Libya/D31 TRACH/2006** | 5.9 | 4.1 | 5.1 | 5.3 | 5.1 | 5.2 | 9.0 | 4.5 | 7.5 |
| **A/ck/Nepal/1-220/2013** | 3.0 | 2.7 | 4.0 | 4.2 | 3.9 | 4.0 | 6.5 | 2.6 | 5.1 |
| **A/ck/Nepal/PT22/2013** | 4.1 | 0.7 | 1.4 | 1.6 | 1.4 | 1.5 | 5.7 | 5.6 | 7.7 |
| **A/ck/Pakistan/47/2003** | 4.2 | 0.8 | 1.2 | 1.3 | 1.2 | 1.3 | 5.6 | 5.8 | 7.9 |
| **A/ck/Pakistan/UDL 7/2008** | 3.3 | 0.8 | 2.1 | 2.3 | 2.1 | 2.2 | 5.7 | 4.6 | 6.7 |
| **A/ck/Saudi Arabia/A/2010** | 4.3 | 1.1 | 1.9 | 2.1 | 2.0 | 2.0 | 6.2 | 5.4 | 7.7 |
| **A/ck/Saudi Arabia/S11A/2003** | 1.9 | 3.0 | 4.2 | 4.5 | 4.2 | 4.2 | 5.8 | 2.4 | 4.2 |
| **A/ck/Saudi Arabia/3489V08-44/2006** | 4.1 | 0.7 | 1.4 | 1.6 | 1.5 | 1.6 | 5.7 | 5.5 | 7.7 |
| **A/ck/Saudi Arabia/3489V08-47/2007** | 4.8 | 2.3 | 3.4 | 3.5 | 3.4 | 3.5 | 7.4 | 4.7 | 7.4 |
| **A/ck/Saudi Arabia/3489V08-50/2008** | 4.0 | 0.6 | 1.3 | 1.5 | 1.3 | 1.4 | 5.5 | 5.6 | 7.7 |
| **A/ck/Saudi Arabia/3489V08/2005** | 4.5 | 2.2 | 3.3 | 3.4 | 3.3 | 3.4 | 7.2 | 4.4 | 7.1 |
| **A/ck/Shijiazhuang/2/1999** | 3.8 | 4.4 | 5.7 | 6.0 | 5.7 | 5.8 | 7.8 | 1.0 | 4.0 |
| **A/ck/Sichuan/5/1997** | 5.1 | 4.4 | 5.7 | 5.9 | 5.7 | 5.8 | 8.8 | 2.8 | 5.8 |
| **A/ck/Tunisia/12/2010** | 3.5 | 1.1 | 2.2 | 2.4 | 2.4 | 2.3 | 5.6 | 6.9 | 8.2 |
| **A/ck/Tunisia/812/2012** | 3.7 | 1 | 2.2 | 2.4 | 2.2 | 2.3 | 6.0 | 4.7 | 7.0 |
| **A/ck/United Arab Emirates/H4TR/2011** | 5.7 | 3.3 | 4.3 | 4.4 | 4.3 | 4.4 | 8.4 | 5.0 | 7.9 |
| **A/ma/Finland/LI13384/2010c** | 2.0 | 4.9 | 5.9 | 6.2 | 5.9 | 5.9 | 5.9 | 2.7 | 2.4 |
| **A/dk/Hong Kong/702/1979** | 5.3 | 6.0 | 7.3 | 7.6 | 7.4 | 7.4 | 9.4 | 1.4 | 4.0 |
| **A/dk/Hong Kong/Y280/1997** | 3.5 | 0.7 | 0.7 | 1.0 | 0.7 | 0.7 | 4.5 | 5.8 | 7.5 |
| **A/dk/Hunan/1/2006** | 5.1 | 2.4 | 1.1 | 0.8 | 1.1 | 1.1 | 4.9 | 7.6 | 9.4 |
| **A/ph/Republic of Ireland/PV18/1997** | 3.7 | 5.2 | 6.5 | 6.7 | 6.5 | 6.5 | 7.8 | 0.3 | 2.8 |
| **A/qa/Arkansas/29209/1993** | 2.8 | 4.8 | 5.9 | 6.2 | 5.9 | 5.9 | 6.8 | 1.3 | 2.5 |
| **A/qa/Hong Kong/A28945/88** | 3.7 | 4.6 | 4.9 | 5.1 | 4.8 | 4.8 | 4.9 | 2.4 | 3.6 |
| **A/qa/Hong Kong/G1/1997** | 0.7 | 3.7 | 4.5 | 4.8 | 4.5 | 4.5 | 4.8 | 3.3 | 3.8 |
| **A/qa/United Arab Emirates/302/2001** | 2.1 | 2.1 | 3.3 | 3.6 | 3.3 | 3.3 | 5.5 | 3.3 | 5.1 |
| **A/rt/New Jersey/AI11-1946/2011** | 3.6 | 4.4 | 5.7 | 5.9 | 5.7 | 5.7 | 7.6 | 0.9 | 3.8 |
| **A/sh/Delaware/277/1999** | 3.7 | 5.2 | 6.5 | 6.7 | 6.5 | 6.5 | 7.8 | 0.5 | 2.6 |
| **A/sh/Delaware/9/1996** | 3.8 | 6.3 | 7.4 | 7.7 | 7.4 | 7.4 | 7.7 | 2.1 | 1.0 |
| **A/ty/Israel/1266/2003** | 3.6 | 2.3 | 3.5 | 3.8 | 3.5 | 3.6 | 6.7 | 3.4 | 5.9 |
| **A/ty/Netherlands/11015452/2012** | 2.5 | 5.7 | 6.6 | 6.9 | 6.6 | 6.6 | 5.7 | 3.7 | 2.4 |
| **A/tk/USA/6707-1/1996** | 3.1 | 5.1 | 6.3 | 6.6 | 6.3 | 6.3 | 7.2 | 1.2 | 2.2 |

### Analysis of antigenic cluster transitions

To better define the amino acid signatures involved in the antigenic profile of H9 HA antigens the differences among the prototypic WF10 h9.4.1 (cyan), the h9.4.2 (blue), and the h9.3.9 consensus viruses were further analyzed. The HAU distance between WF10 h9.4.1 and h9.4.2 are lower (3.4 AU) than the distance between WF10 h9.4.1 and h9.3.9 (4.1 AU). Amino acid substitution differences between WF10 h9.4.1 (cyan) and the h9.4.2 (blue) include E72G, G135D, E180A, and I186T, which have been previously reported as antigenically relevant for H9 (12, 13, 16, 17, 34). We selected 9 positions, 72, 131, 135, 150, 180, 186, 188, 198, and 217 that differed between WF10 h9.4.1 and h9.4.2 and changed specific amino acid positions by site-directed mutagenesis. **(Fig 3A-B and Table 4)**. The WF10-9p-h9.4.2 virus expressing the WF10 HA with the 9 amino acid signatures of the h9.4.2 consensus showed antigenic cluster transition from cyan (WF10 h9.4.1) to blue (h9.4.2) (**Fig 3C**). The distance between WF10 h9.4.1 and WF10-9p-h9.4.2 was 3.8 AU, whereas the distance between h9.4.2 and WF10-9p-h9.4.2 was 1.6 AU. The counterpart h9.4.2-9p-WF10 virus expressing the h9.4.2 HA1 portion with the 9 amino acids from WF10 **(Fig 3B)** showed antigen transition from the blue (h9.4.2) to cyan (WF10 h9.4.1) cluster (**Fig 3C)**. The distance between h9.4.2 and h9.4.2-9p-WF10 was 3.2 AU, whereas the distance between h9.4.2-9p-WF10 and WF10 h9.4.1 was 0.7 AU confirming the antigenic relevance of these positions. Similarly, two WF10 h9.4.1 viruses **(Fig 4A-B)** carrying 7 amino acid signatures of h9.3.9 (WF10-7p-h9.3.9a: 127, 131, *173*, 180, 182, 183, and 217 and WF10-7p-h9.3.9b: 127, 131, *146*, 180, 182, 183, and 217) showed full cluster transition from cyan (h9.4.1) to orange (h9.3.9) (**Fig 4D and Table 4**) with 0.9 AU and 1 AU of distance between h9.3.9 and WF10-7p-h9.3.9a or WF10-7p-h9.3.9b, respectively. The h9.3.9-8p-WF10 virus with 8 amino acid signatures positions (127, 131, 146, 173, 180, 182, 183, and 217) of the WF10 h9.4.1 (**Fig 4C**) showed antigenic transition from orange (h9.3.9) to cyan (WF10 h9.4.1) (**Fig 4D**). Distances between h9.3.9-8p-WF10 (cyan) and h9.3.9 (orange) or WF10 h9.4.1 (cyan) were 3.6 AU and 1.3 AU, respectively. To further characterize antigenically relevant amino acid positions in more detail, single and double mutants in the context of WF10 h9.4.1 were produced **(Figs 5-6 and Table 5)**. From a panel of 19 mutants produced, 14 were viable. The results showed that the E180A-h9.4.2 single mutant **(Fig 5C)** and the R131K/E180A-h9.4.2 double mutant **(Fig 5E)** led to the most significant antigenic changes between WF10 h9.4.1 (cyan) and h9.4.2 (blue). In both cases, antigens were cross-reactive between the cyan and blue clusters, determined by an AU <3 from WF10 h9.4.1 (cyan) and h9.4.2 (blue). The remaining single and double mutants affected HI activity **(Tables 4 and 5)**, but none resulted in cluster transitions. Taken together, the results show that different positions modulate with different magnitudes the antigenic properties of H9 HA. Amino acid 180 has, in general, the largest effect on HI activity.

**Fig. 3.**
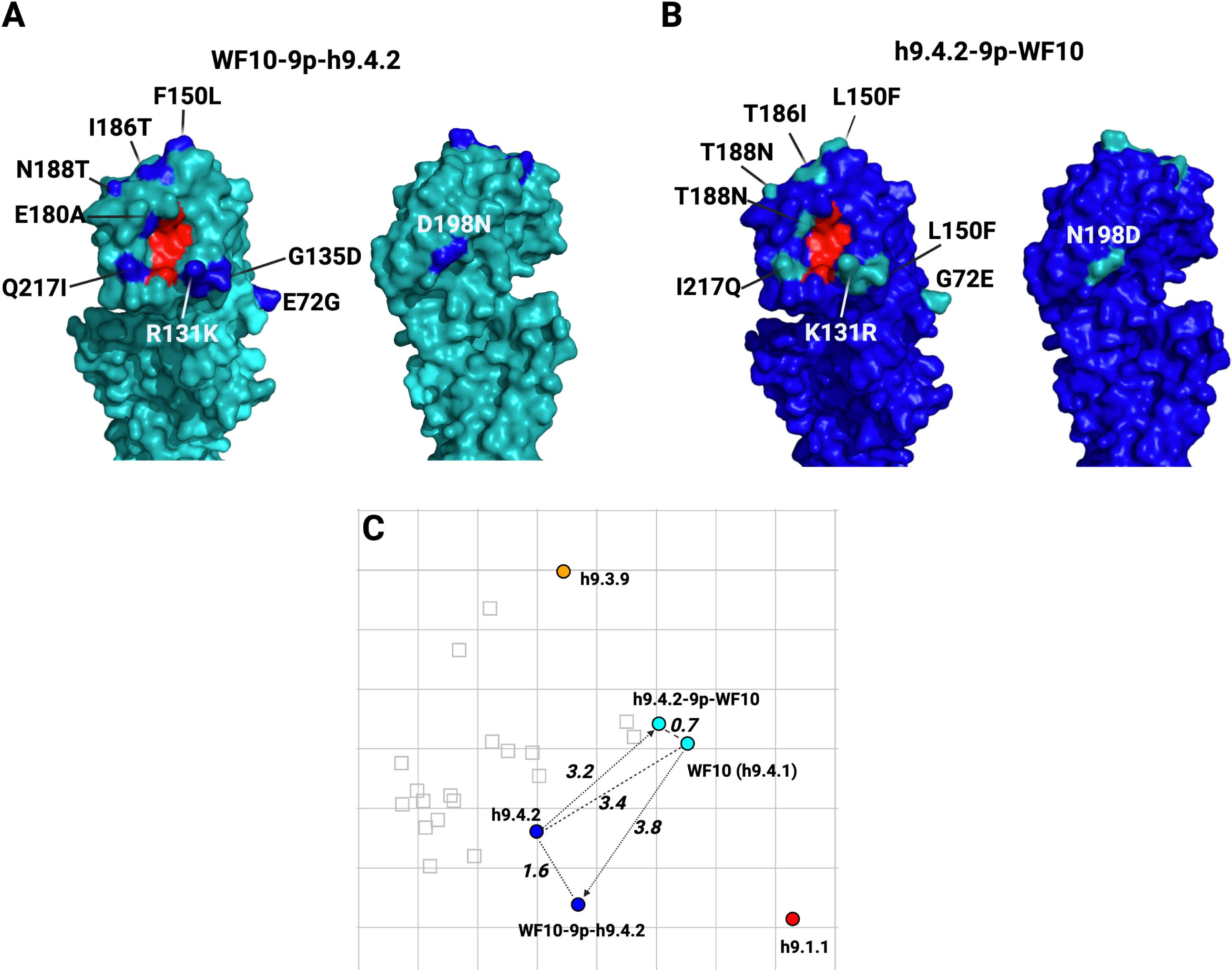
Analysis of molecular signatures of antigenicity between cyan-blue antigenic clusters. Transitions of H9 virus mutants carrying selected amino acid substitutions. 3D structures were generated with PyMOL and color-coded as follows: Red = amino acids in the RBS (91, 143, 173, 184, 185, and 216), cyan=WF10, and blue= h9.4.2. (**A**) WF10-9p-h9.4.2 mutant with the HA-WF10 carrying substitutions at positions 72, 131, 135, 150, 180, 186, 188, 198, and 217 corresponding to the h9.4.2 consensus sequence. **(B)** h9.4.2-9p-WF10 mutant with the HA-h9.4.2 modified at amino acid positions 72, 131, 135, 150, 180, 186, 188, 198, and 217 corresponding to the WF10 HA sequence. **(C)** Antigenic map showing the antigenic cluster transitions of the mutants evaluated. AUs are stated adjacent to respective arrows: Dashed lines highlight the distances between the mutant and the “target” virus and between WF10 (h9.4.1) and h9.4.2.

**Fig. 4.**
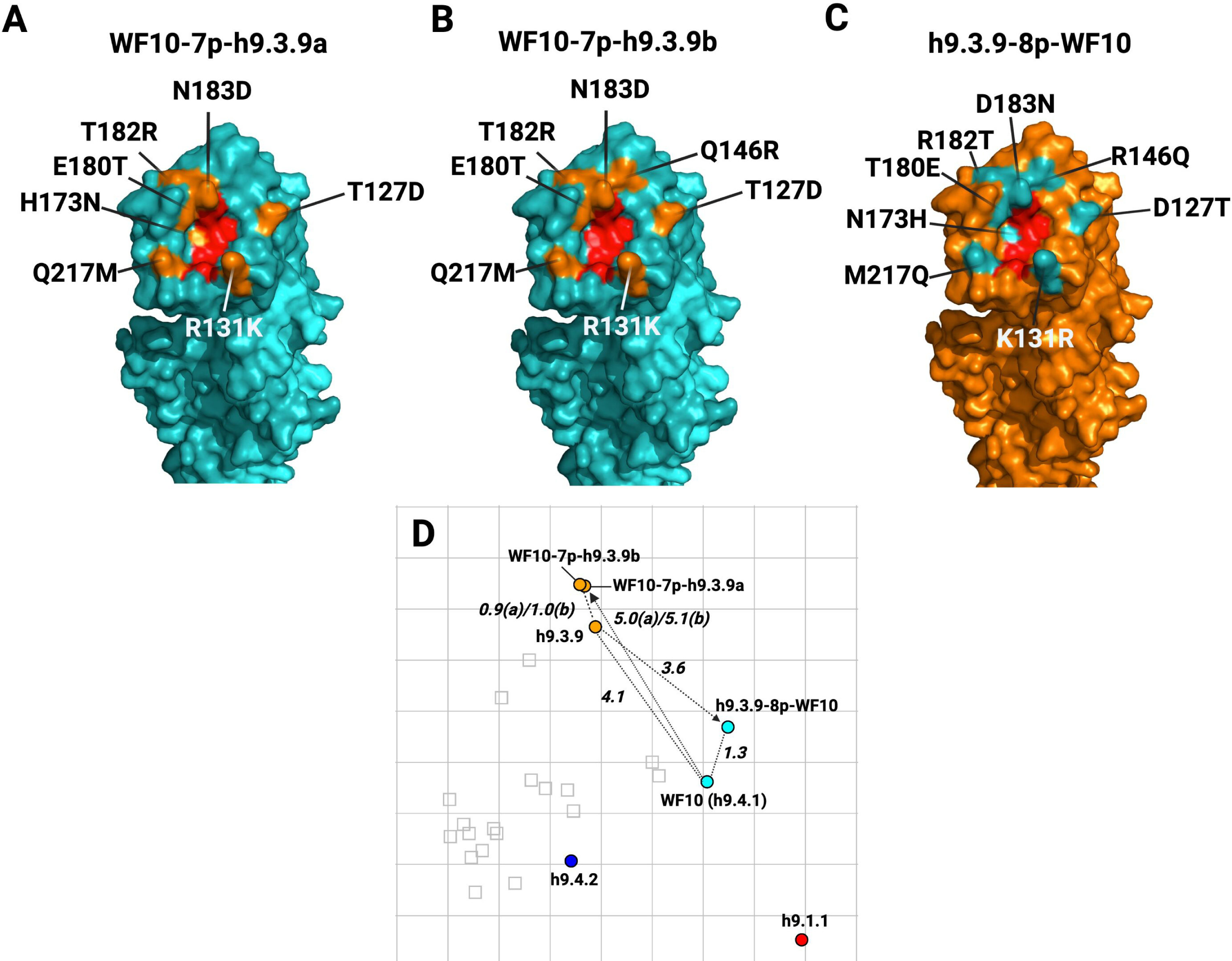
Cyan-orange antigenic cluster transitions of H9 virus mutants carrying selected amino acid substitutions. **(A)** WF10-7p-h9.3.9a mutant with the HA-WF10 H9.4.1 and substitutions at positions 127, 131, ***173***, 180, 182, 183, and 217 corresponding to the h9.3.9 antigen. **(B)** The WF10-7p-h9.3.9b mutant is the same as in A, except that substitutions are at positions 127, 131, ***146***, 180, 182, 183, and 217. **(C)** h9.3.9-8p-WF10 mutant with the HA-h9.3.9 modified at amino acid positions 127, 131, 146, 173, 180, 182, 183, and 217, corresponding to the WF10 HA sequence. 3D structures as described in **Fig 3**, except that orange, highlights amino acids in the h9.3.9 consensus sequence. **(D)** Antigenic map showing the antigenic cluster transitions of the mutants evaluated. AUs are stated adjacent to respective arrows: Dashed lines highlight the distances between the mutant and the “target” viruses and between WF10 (h9.4.1) and h9.3.9.

**Fig. 5.**
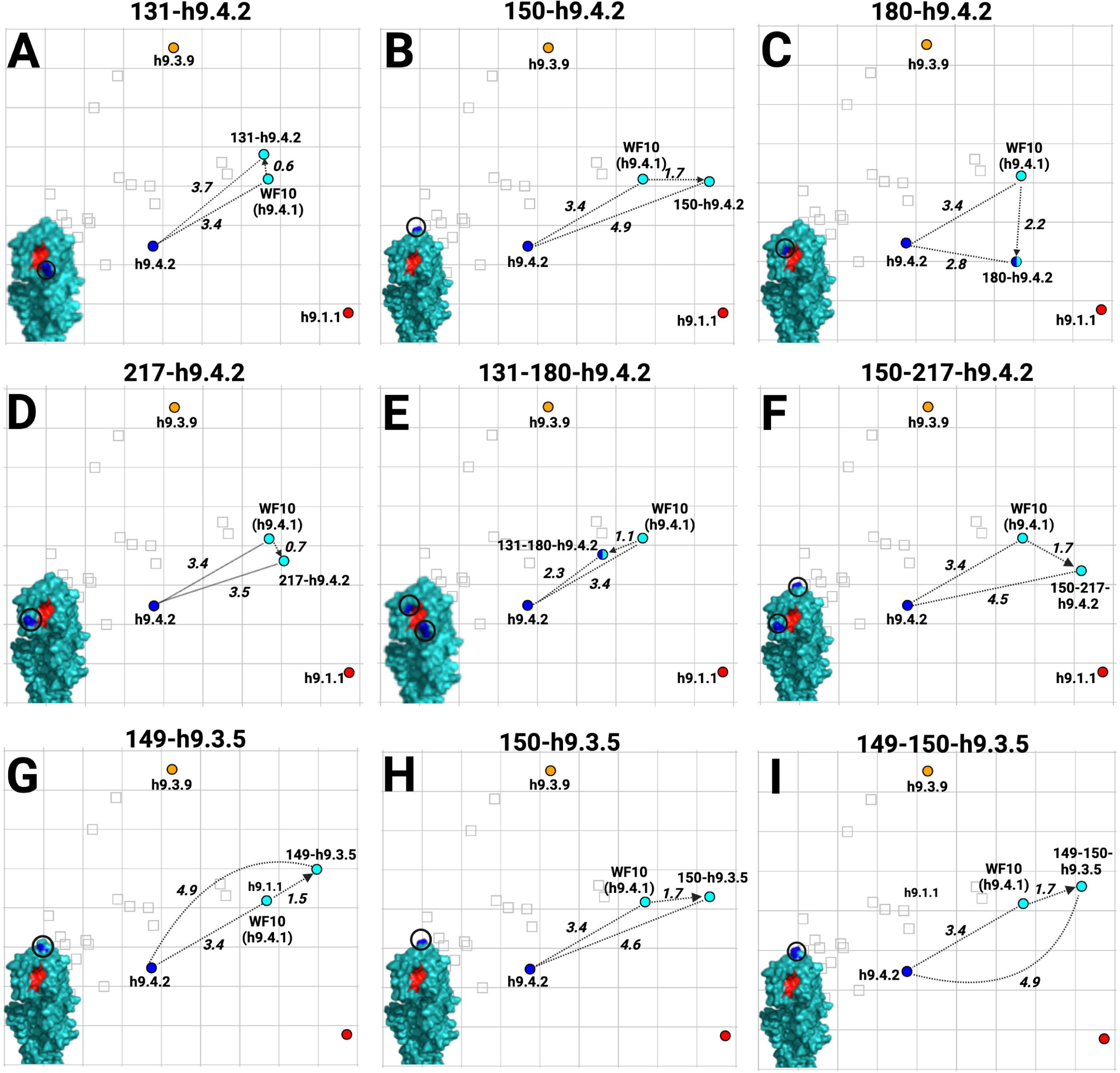
Antigenic cartography results for H9 virus mutants carrying single or double amino acid substitutions between WF10 h9.4.1 and h9.4.2/h9.3.5 in the WF10 HA backbone. **(A)** R131K-h9.4.2; **(B)** F150L-h9.4.2; **(C)** E180A-h9.4.2 **(D)** Q217I-h9.4.2; **(E)** R131K-E180A-h9.4.2; **(F)** F150L-Q217I-h9.4.2; **(G)** G149N-h9.3.5; **(H)** F150A-h9.3.5; **(I)** G149N-F150A-h9.3.5. AUs and 3D renderings are color-coded as described in **Fig. 3**. Only the E180A-h9.4.2 (cyan to blue) and the R131K/E180A-h9.4.2 (cyan to blue) mutants showed cluster transitions.

**Fig. 6.**
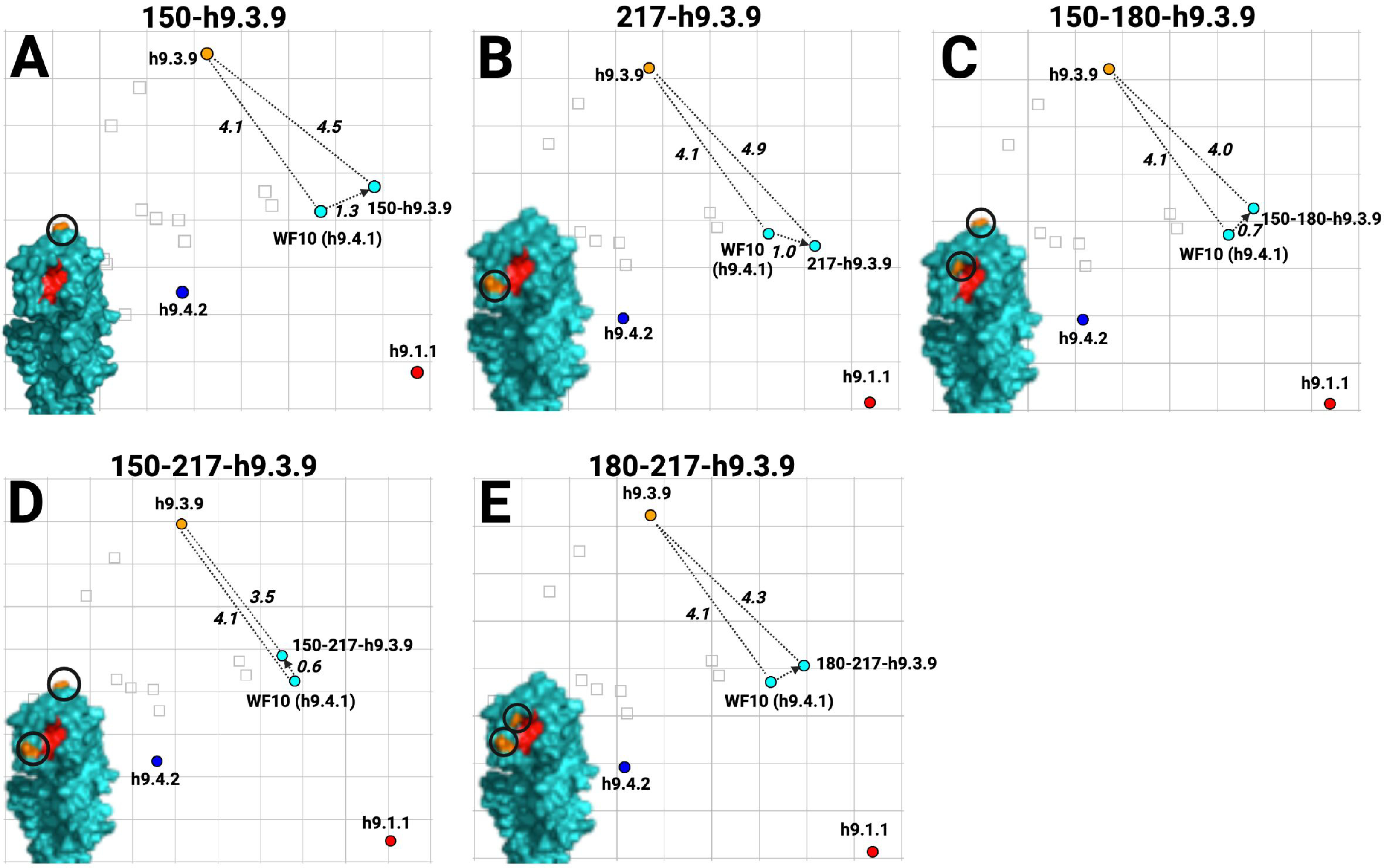
Antigenic cartography results for H9 virus mutants carrying single or double amino acid substitutions between WF10 h9.4.1 and h9.3.9 in the WF10 HA backbone. **(A)** F150D-h9.3.9; **(B)** Q217M-h9.3.9; **(C)** F150D-E180T-h9.3.9; **(D)** F150D-Q217M-h9.3.9; **(E)** E180T-Q217M-h9.3.9. AU units and 3D renderings are color-coded, as described in **Fig. 4**.

**Table 4.** Summary of amino acid substitutions for each mutant and antigenic distances

| Mutant | Substitution | Virus | Antigenic distance from |  | Antigenic cluster |
| --- | --- | --- | --- | --- | --- |
|  |  |  | WF10 | h9.4.2 |  |
| WF10 (H9N2) | Wild type | YES | 0.0 | 3.4 | *cyan* |
| h9.4.2 | HA1 | YES | 3.4 | 0.0 | *blue* |
| WF10-9p-h9.4.2 | E72G, R131K, G135D, F150L, E180A, I186T, N188T, D198N, Q217I | YES | 3.8 | 1.6 | *blue* |
| h9.4.2-9p-WF10 | G72E, K131R, D135G, L150F, A180E, T186I, T188N, N198D, I217Q | YES | 0.7 | 3.2 | *cyan* |
| 131-h9.4.2 | R131K | YES | 0.6 | 3.7 | *cyan* |
| 150-h9.4.2 | F150L | YES | 1.7 | 4.9 | *cyan* |
| 180-h9.4.2 | E180A | YES | 2.2 | 2.8 | *blue/cyan* |
| 217-h9.4.2 | Q217I | YES | 0.7 | 3.5 | *cyan* |
| 131-180-h9.4.2 | R131K/E180A | YES | 1.1 | 2.3 | *blue/cyan* |
| 150-180-h9.4.2 | F150L/E180A | NO | - | - | - |
| 150-217-h9.4.2 | F150L/Q217I | YES | 1.7 | 4.5 | *cyan* |
| 180-217-h9.4.2 | E180A/Q217I | NO | - | - | - |
|  | **Substitution** | **Virus** | **WF10** | **h9.3.9** | **Antigenic cluster** |
| h9.3.9 | HA1 | YES | 4.1 | 0.0 | *orange* |
| WF10-7p-h9.3.9a | T127D, R131K, V173M, E180T, T182R, N183D, Q217M | YES | 5.0 | 0.9 | *orange* |
| WF10-7p-h9.3.9b | T127D, R131K, Q146R, E180T, T182R, N183D, Q217M | YES | 5.1 | 1.0 | *orange* |
| h9.3.9-8p-WF10 | D127T, K131R, R146Q, M173V, T180E, R182T, D183N, M217Q | YES | 1.3 | 3.6 | *cyan* |
| 150-h9.3.9 | F150D | YES | 1.3 | 4.5 | *cyan* |
| 180-h9.3.9 | E180T | NO | - | - | - |
| 217-h9.3.9 | Q217M | YES | 1.0 | 4.9 | *cyan* |
| 150-180-h9.3.9 | F150D/E180T | YES | 0.7 | 4.0 | *cyan* |
| 150-217-h9.3.9 | F150D/Q217M | YES | 0.6 | 3.5 | *cyan* |
| 180-217-h9.3.9 | E180T/Q217M | YES | 0.7 | 4.3 | *cyan* |
|  | **Substitution** | **Virus** | **WF10** | **h9.3.5** | **Antigenic cluster** |
| h9.3.5 | HA1 | YES | 4.1 | 0.0 | *blue* |
| 149-h9.3.5 | G149N | YES | 1.5 | 5.6 | ***cyan*** |
| 150-h9.3.5 | F150A | YES | 1.7 | 5.7 | ***cyan*** |
| 180-h9.3.5 |  | YES | 1.3 | 4.2 | ***cyan*** |
| 149-150-h9.3.5 | G149N/F150A | YES | 1.5 | 5.6 | ***cyan*** |

**Table 5.** Antigenic distances between consensus viruses and the different mutants.

|  | WF10 | h9.4.2 | h9.3.3 | h9.3.4 | h9.3.5 | h9.3.7 | h9.3.9 | h9.1.1 | h9.2.2 |
| --- | --- | --- | --- | --- | --- | --- | --- | --- | --- |
| 131-180-4.2 | 1.1 | 2.3 | 3.0 | 3.3 | 3.0 | 3.0 | 4.0 | 4.3 | 5.3 |
| 131-4.2 | 0.6 | 3.7 | 4.2 | 4.5 | 4.2 | 4.2 | 3.6 | 4.6 | 4.8 |
| 149-150-H9.3.5 | 1.5 | 4.9 | 5.7 | 5.9 | 5.6 | 5.6 | 4.8 | 3.9 | 3.3 |
| 149-H9.3.5 | 1.5 | 4.9 | 5.6 | 5.8 | 5.6 | 5.5 | 4.5 | 4.3 | 3.7 |
| 150-180-3.9 | 0.7 | 4.1 | 4.8 | 5.0 | 4.7 | 4.7 | 4.0 | 4.2 | 4.2 |
| 150-217-3.9 | 0.6 | 3.5 | 4.0 | 4.3 | 4.0 | 4.0 | 3.5 | 4.6 | 4.9 |
| 150-217-4.2 | 1.7 | 4.5 | 5.5 | 5.8 | 5.5 | 5.5 | 5.7 | 2.6 | 2.8 |
| 150-H9.3.5 | 1.7 | 4.9 | 5.8 | 6.0 | 5.7 | 5.7 | 5.2 | 3.6 | 3.0 |
| 150-H9.3.9 | 1.3 | 4.6 | 5.4 | 5.6 | 5.3 | 5.3 | 4.5 | 4.0 | 3.6 |
| 150-H9.4.2 | 1.7 | 4.9 | 5.8 | 6.0 | 5.8 | 5.7 | 5.3 | 3.4 | 2.9 |
| 180-217-3.9 | 0.7 | 4.1 | 4.8 | 5.0 | 5.1 | 4.8 | 4.8 | 4.3 | 4.0 |
| 180-H9.3.5 | 1.3 | 3.2 | 4.2 | 4.5 | 4.2 | 4.2 | 5.3 | 2.8 | 4.0 |
| 180-H9.4.2 | 2.2 | 2.8 | 4.1 | 4.3 | 4.1 | 4.1 | 6.0 | 2.5 | 4.5 |
| 217-3.9 | 1.0 | 4.1 | 5.0 | 5.3 | 5.0 | 5.0 | 4.9 | 3.3 | 3.5 |
| 217-4.2 | 0.7 | 3.5 | 4.4 | 4.7 | 4.4 | 4.4 | 4.8 | 3.3 | 3.9 |
| 3.9-8PWF10 | 1.3 | 4.5 | 5.0 | 5.2 | 4.9 | 4.9 | 3.6 | 4.8 | 4.5 |
| 4.2-9PWF10 | 0.7 | 3.2 | 3.7 | 4.0 | 3.7 | 3.6 | 3.5 | 4.6 | 5.1 |
| WF10-7P-3.9 C3 | 5.0 | 5.9 | 5.4 | 5.4 | 5.4 | 5.3 | 0.9 | 8.9 | 8.8 |
| WF10-7P-3.9 C4 | 5.1 | 6.0 | 5.4 | 5.4 | 5.4 | 5.3 | 1.0 | 9.0 | 8.9 |
| WF10-9P-H9.4.2 | 3.8 | 1.6 | 2.8 | 3.0 | 2.9 | 2.9 | 6.5 | 4.2 | 6.6 |

## DISCUSSION

The HA has a pivotal role in the antigenicity of FLUAV as it is the major target of neutralizing antibodies and subject to positive selection. Phylogenetics combined with antigenic analysis is the basis for human, avian, and equine influenza vaccine selection (43). Antigenic cartography facilitates the understanding of FLUAV antigenic drift by visualizing HI data as a spatial relationship between antigens in a map (36, 38, 42). We captured the antigenic diversity of dissimilar H9 viruses, underscored by the ability of synthetic consensus viruses to induce HI responses that recognize their genetically related field antigens. For antigenic characterization, boost immunizations with inactivated-whole virus adjuvant formulations were performed in quail at 14 dpi and allowed increasing titer levels of poorly immunogenic antigens **(Fig. 2A)**. Quail antibody responses to H9 FLUAV mimicked what was previously reported in the literature for chicken sera (13, 17). The synthetic consensus viruses aligned antigenically with representative H9 prototype field strains, supporting that the HA globular head has a pivotal role in shaping the antigenic phenotype. The results reinforce the hypothesis that genetic relatedness can predict the antigenic phenotype, with some exceptions (**Fig 2B-D**). WF10 h9.4.1 and A/qa/HK/G1/97 (G1 prototype strain), which are phylogenetically related, showed also antigenic similarity (cyan cluster). These two antigens clustered separately from h9.4.2 (blue cluster), which showed strong cross-reactivity with most poultry isolates from the Middle East and Asia **(Table 2)**. Similarly, h9.3.3 and h9.3.4 consensus antigens demonstrated strong cross-reactivity with their respective prototype lineages A/ck/HK/G9/1997 (G9, h9.3.3-like) and A/dk/HK/Y280/1997 (Y280, h9.3.4-like). The strong HI cross-reactivity of the H9 field isolates against the heterologous clade-specific consensus antisera also supported the antigenic map results. Interestingly, consensus clades h9.3.3-7 and h9.4.2 showed similar antigenic phenotypes despite their genetic differences. Furthermore, the h9.3.9 (orange cluster) antigenic properties differed significantly from the rest of the h9.3 consensus viruses, with the highest reaction against its homologous sera (HI titer: 5120) and marginal cross-reactivity with heterologous sera. Strikingly, the % of identity between the h9.3.9 consensus HA and the closest relative (A/dk/Hunan/1/2006) was 93.3%, being the lowest observed among the different clades, perhaps exposing a gap in sequence availability from the online databases. Nonetheless, sequence comparison between h9.3.9 and A/dk/Hunan/1/2006 revealed differences in key positions such as G72E, R146Q, N149G, N183D, and M217Q **(Fig 4-6)**(12, 13, 16, 17, 34) which may account for the antigenic differences despite the close phylogenetic relationships. Few other H9 field isolates fell outside the 3 AU radius from any consensus antigen, despite the intermediate level of reactivity against the antisera panel. These observations highlight the significant impact of a few amino acid changes in modulating HI activity (35-38) and reiterate the importance of antigenic cartography in correcting phylogenetic predictions.

Most wild bird isolates from Europe and North America clustered with the h9.1.1 and h9.2.2 consensus antigens (red cluster) **(Table 2),** as predicted by the phylogenetic analysis. However, this data must be carefully interpreted due to the relatively low inherent antigenicity and immunogenicity of these antigens compared to the rest of the consensus and field antigens. This was evidenced by the relatively low homologous HI titers (40-160) obtained in quail immunized with h9.1.1 (HI titers: 80-160) and h9.2.2 (HI titers: 40-160) despite the boost and the overall poor cross-reactivity of these antigens with heterologous sera **(Table 1)**. Similarly, most H9 wild bird isolates from Europe and North America had poor reactivity with any consensus heterologous sera **(Table S1)**.

The generation of a humoral response that interferes with the interaction of HA with its receptor is key to achieving sterilizing immunity against FLUAV. Seven residues (145, 155, 156, 158, 159, 189, and 193, H3 numbering) near the RBS were identified as the major determinants of antigenic drift in human and swine H3N2 FLUAVs (35, 39). Similarly, amino acid substitutions were identified as the major drivers of antigenic diversity of H5N1 clade 2.1, human H2N2, and pandemic H1N1 FLUAVs (37, 40, 41). For H9N2, molecular signatures of antigenicity are poorly characterized. Over 40 amino acid positions have been described for the H9 HA as antigenically relevant, mainly through generating escape mutants using mouse monoclonal antibodies and/or inferred from HI data (13, 14, 16, 19, 42-45). Using chicken sera, 24 amino acid positions distributed over the entire H9 HA were considered antigenically relevant (17). Based on the initial antigenic characterization **(Fig 2),** full cluster transitions from WF10 (cyan) to the h9.4.2 (blue) and h9.3.9 (orange) antigenic profiles were readily observed with the WF10-9p-h9.4.2 (substitutions at positions 72, 131, 135, 180, 186, 188, 198, and 217) and WF10-7p-h9.3.9a/b antigens (substitutions at positions 127, 131, 146 or 173, 180, 182, 183, and 217), respectively **(Fig 3 and 4)**. The impact of single or double amino acid substitutions was less clear **(Figs 5 and 6)**. The E180A-h9.4.2 single mutant **(Fig 5C)** and the R131K/E180A-h9.4.2 double mutant **(Fig 5E)** showed the strongest effect, with antigens positioning at <3.0 AU from the cyan and blue antigenic cluster. These observations suggest a role for position 180 since the R131K single mutant had minimal effect on HI activity compared to the WF10 h9.4.1 HA **(Fig 5A)**. Consistent with these observations, a previous report using the strain A/chicken/Shanghai/F/98 (H9N2) determined position 180 as directly responsible for antigenic drift (30). Variability at position 180 was also reported in field isolates from Morocco between 2018-2019, reinforcing a preponderant role of position 180 in evading pre-existing immunity (46). Consistent with the Morocco study, molecular characterization of H9N2 viruses from local markets in southern China also revealed a potential role of position 180 (and other positions) on antigenic properties (28). Spatiotemporal dynamics analysis from live-poultry markets in China has shown selection pressure in positions 146, 150, and 180 (47). A role of position 180 has been suggested also for the cross-species barrier where the 180V mutation favors the replication of H9N2 in mice (48). Other studies have attributed antigenic modulation to several HA residues without including position 180 (27, 49). The latter is consistent with the idea that additional positions within the HA can modulate the antigenic properties, which is consistent with the findings in this report where 8 or 9 substitutions were introduced **(Fig 3 and 4)**. A previous report also described the role of position 217 in H9 antigenicity. However, in the global scale analysis, position 217 alone is insufficient for an antigenic cluster transition suggesting modest effects on antigenicity (29). Position 183 was also recently suggested as a modulator of the antigenic properties and overall replication of H9N2 viruses (50). This is consistent with the results observed between WF10 h9.4.1 (cyan) and h9.4.2 (blue). Antigenically relevant positions such as 180 and 217 have also been shown to affect receptor-binding avidity (29, 51, 52), as it has position 216 (4, 24, 53).

Other single or double substitutions showed changes in the level of antigenicity; however, none were enough on their own to produce complete antigenic cluster transitions (**Fig. 5**). Reduced HI activity against the parental WF10 antiserum was observed for substitutions at positions G149N-h9.3.5 and F150A-h9.3.5 (1.7 AU and 1.5 AU respectively), but no reciprocal increase in cross-reactivity against the target antiserum was observed **(Table 4)**. The F150L/Q217I-h9.4.2 double mutant had a higher impact on the parental WF10 than the single mutant Q217I-h9.4.2 (1.7 AU for 150-217-h9.4.2 versus 0.7 AU for 217-h9.4.2), and a similar effect was observed against the target h9.4.2 antiserum (4.5 AU for F150L/Q217I-h9.4.2 versus 3.5 AU for Q217I-h9.4.2). These observations point to relatively few additional substitutions as likely responsible for antigenic cluster transitions.

Despite the remarkable plasticity of the H9 HA of WF10, reversions were observed in 5 out of 19 mutants, suggesting that tolerability of changes in antigenically relevant amino acids may be context dependent and likely encompass compensatory substitutions (36, 54). In addition, we identified a set of non-cross-reactive strains **(Table 2 and Fig 2C)** whose initial sequence information would predict to fall in at least one of the antigenic clusters described. These strains included A/dk/HK/448/1978, A/qa/Saudi A/489_46v08/2006, A/ck/NKorea/99020/99 and A/ma/Eng/7798_6499/2006. The strain, A/ck/Tun/345/2011, with an HA1 region almost identical to A/ck/Tun/812/2012 in key amino acid signatures, failed to show cross-reactivity with members of the blue cluster, suggesting the involvement of other potentially relevant epitopes.

Most studies of H9N2 antigenicity in poultry involve the use of chicken sera but not sera from minor land-based poultry species, such as quail. Japanese quails have been suggested as key players in the genesis of the adaptation of influenza viruses with respiratory tract tropism (21, 22). Quail are also more susceptible to H9N2 infection than chickens (31). In addition, quail show wide distribution in the respiratory tract of avian-like (SAα2.3) and human-like (SAα2.6) sialic acid receptors, which may have contributed to the emergence of the current poultry-adapted H9N2 strains with human-like receptor preference (55). Thus quail might have played a role as an intermediate host between wild aquatic birds and poultry in the emergence of H9N2 strains with altered host range (23, 24). The antigenic analyses using quail antisera provide significant insights into anti-HA responses in a relevant poultry species for influenza replication and evolution. The current literature shows different approaches employed for the antisera generation, including live virus inoculation and inactivated/adjuvanted virus vaccination to study antigenicity of the HA of influenza viruses. Still, none have used quail sera as a model (17, 25-28). The results validate using the quail model to study the antigenicity of H9N2 as well as other viral properties such as virus replication, pathogenesis, and transmission. Although the results provide novel insights into the antigenic properties of FLUAV of the H9 subtype on a global scale, some limitations must be noted. The initial phylogenetic analysis for generating the consensus sequences was performed in 2016. As H9N2 viruses continue to evolve with inherent animal and public health risks, further studies are needed to better dissect the role of amino acid substitutions on the HA that modulate host range, replication, pathogenesis, transmission, and antigenicity.

In conclusion, phylogenetics was used to generate consensus on H9 viruses encompassing their natural diversity. We demonstrated that these consensus H9 viruses were biologically active, capable of triggering an immune response associated with the generation of neutralizing antibodies, and manifested important distinctive biologic characteristics driven only by their differences in the HA1 domains. Using this system, we explored antigenicity and modulation of HI profiles using antisera obtained from quail. The sera obtained allowed us to narrow down antigenically relevant amino acids, as many as 9 for h9.4.2 (at positions 72, 131, 135, 180, 186, 188, 198, and 217) and 6 for h9.3.9 (127, 131, 180, 182, 183, and 217) to as few as 1 (E180A), to produce antigenic cluster transitions. The results are relevant to pave the way for a better understanding of the molecular signatures of antigenicity in H9 viruses, facilitating a rational approach for selecting more efficacious vaccines against poultry-origin H9 influenza viruses.

## MATERIALS AND METHODS

### Ethics statement

Use of quail for sera preparation against H9 FLUAVs adhered to and approved by the Institutional Animal Care and Use Committee of UGA under protocols A201506-026-Y3-A5. Quail studies were conducted in a USDA-approved ABSL2 facility at the Poultry Diagnostic Research Center, College of Veterinary medicine, UGA, with each group of quail housed in individual HEPA in/out isolator units. As needed, based on humane endpoints or at the end of the experiments, animals were humanely euthanized following guidelines approved by the American Veterinary Medical Association.

### Cells

Madin-Darby canine kidney (MDCK) cells were a kind gift from Robert Webster (St. Jude Children’s Research Hospital, Memphis, TN). Human embryonic kidney 293T cells were obtained from the American Type Culture Collection (CRL-3216, Manassas, VA). Cells were maintained in Dulbecco’s modified Eagle’s medium (DMEM; Sigma-Aldrich, St. Louis, MO) containing 10% fetal bovine serum (Sigma-Aldrich), 1% antibiotic-antimycotic (Sigma-Aldrich) and 1% L-glutamine (Sigma-Aldrich). Cells were cultured at 37°C in a humidified incubator under 5% CO2.

### Database and phylogenetic analysis of HA sequences

H9 HA sequences were obtained from the Influenza Research Database (IRD), the Bacterial and Viral Bioinformatics Resource Center (BV-BRC), and the Global Initiative on Sharing All Influenza Data (GISAID) (56, 57). The initial phylogenetic analysis was performed on 984 global representative H9 avian isolates from 1966 to the 18th of March 2016 and was used to build the H9 consensus sequences presented in this study. The phylogenetic analysis was then updated on July 14^th^, 2020, and included 1,316 manually curated sequences. The amino acid frequencies were analyzed using the protein sequence variant analysis tool provided by Scop3D (58). HA sequences were mapped to the A/gf/HK/WF10/1999 (WF10), GenBank accession #AY206676, (31) reference sequence using Geneious (version 10.2.3, Auckland, New Zealand). H9 HA1 sequences spanning the period from 1966 to 2020 were manually pruned to remove truncated and or repetitive sequences. An amino acid alignment was generated using default settings in MUSCLE (59). The numbering of HA corresponds to the mature H9 HA. All known key antigenic sites were considered in the phylogenetic algorithm using optimization with GARLI (60). A maximum-likelihood tree was inferred using RAxML v.8.1.24 (61) with a general time-reversible (GTR) substitution model with gamma-distributed rate variation among sites, followed by Garli for branch optimization. A starting tree was generated using parsimony methods with the best-scoring tree, and statistical support was obtained using the rapid bootstrap algorithm. Initially, 18 consensus sequences were produced, representative of genetic variations within phylogenetic groups. Of these, 10 consensus sequences were selected (**Fig. 1)**.

### Generation of chimeric HA plasmids for reverse genetics

The cDNA copies encoding the HA1 consensus or mutant sequences were synthesized by Genscript (Piscataway, NJ, USA) and then sub-cloned into the plasmid pDP-BsmbI-WF10_HA2 encoding the HA2 portion of WF10. All chimeric constructs contained the identical cleavage site motif (PARSSR) of the WT HA of WF10.

### Viruses

Chimeric HA plasmids were used for reverse genetics using the previously described WF10 backbone (53, 62). Reverse genetics was performed using co-cultured 293T and MDCK cells, as previously detailed (63). Virus stocks were prepared in MDCK cells or 9 to 11-day-old specific pathogen-free (SPF) embryonated chicken eggs. Virus stocks were aliquoted and stored at −80°C until use. Virus stocks were titrated by tissue culture infectious dose 50 (TCID_50_) as described (64).

### Sequencing

Standard Sanger sequencing was performed on all HA plasmids and HA PCR products from all H9 virus stocks by Psomagen (Rockville, MD, USA). Next-generation sequencing (NGS) was performed on all consensus viruses’ whole genomes to exclude unwanted substitutions. For whole-genome sequencing, amplicon sequence libraries were prepared using the Nextera XT DNA library preparation kit (Illumina, San Diego, CA) according to the manufacturer’s protocol. Barcoded libraries were multiplexed and sequenced on a high-throughput Illumina MiSeq sequencing platform in a paired-end 150-nucleotide run format. De novo genome assembly was performed as described previously (65).

### Preparation of H9 antisera in quail

3-week-old Japanese quails (*Coturnix c. japonica,* n=6/group) were inoculated by the oculo-nasal-tracheal route with 10^6^ TCID50/quail of the following WF10-chimeric HA (H9N2) viruses: h9.1.1, h9.2.2, h9.3.3, h9.3.4, h9.3.5, h9.3.7, h9.3.9, h9.4.1 (WF10) and h9.4.2. A negative control (n=6, mock-inoculated with PBS) was included. Active infections were monitored by Flu DETECT (Zoetis, Kalamazoo, Michigan) on tracheal swabs collected from days 1-7 post-inoculation. Boost vaccination was performed with the homologous virus inactivated at 4°C for 3 days with 0.1% beta propiolactone (BPL) (Sigma-Aldrich Corporation, St. Louis, MO) as previously described (66). On the day of the boost, 512-1024 HAU/50ul of the corresponding virus was mixed 1:1 (vol/vol) with Montanide ISA 71 VG adjuvant (Seppic, Paris, France), in an emulsion, as per manufacturer protocol. Then, quail were inoculated subcutaneously in the neck with 300 µL (150 µL inactivated virus + 150 µL Montanide) of the homologous virus-adjuvant emulsion. At 14 days post-boost (dpb), quails were terminally bled under anesthesia, and sera were collected for HI assays. After testing each bird’s seroconversion level, sera with similar titers were pooled, three quail/pool, two sera pools/antigen (**Table 1**).

### Antigenic characterization

Standard hemagglutination (HA) and HI assays were performed as previously described (67). Before HI testing, sera were heat inactivated at 56°C for 30 min and adsorbed with 50% chicken red blood cells (RBCs) to remove nonspecific inhibitors of hemagglutination. Sterile PBS was added, allowing the sera to reach a final dilution of 1:10. Then sera were transferred to 96-well plates and serially diluted 2-fold in 25 µL of sterile PBS and mixed with 4 HAU/25 µL of each virus. The virus-sera mixture was incubated for 15 min at room temperature and then added 50 µL per well of 0.5% chicken RBCs (100 µL final volume/well). The HI activity was determined after 45 min of incubation.

### Antigenic cartography

The HI data using quail sera **(Table S1)** was analyzed separately and merged through the ACMACS antigenic cartography website (https://acmacs-web.antigenic-cartography.org) as previously described (68, 69). HI data sets were subject to a dimensional analysis in all dimensions (2D, 3D, 4D and 5D) with 2,000 optimizations and an automatic minimum column basis parameter to identify which model best fits this data set. Antigens that exploited no to low (<40) reactivity against the entire antisera panel were removed from the analysis and annotated. The distance between the spheres (antigens) and antisera (squares) is inversely correlated to the log_2_ titer measured by the HI assay. One antigenic unit is the equivalent of a 2-fold loss/gain in HI activity. Clusters were initially established by applying the Ward method of hierarchical clustering. Within these, reference antigens were selected based on their biological significance, and clusters were adjusted to enclose antigens exclusively within a 3 AU radius from these selected reference antigens. We used 3 AU or a ≥8-fold loss in cross-reactivity, as defined by the WHO recommendation to update human seasonal vaccine strains, as the threshold of significant antigenic difference.

### Site Direct Mutagenesis

The site-directed mutagenesis kit (ThermoFisher, Waltham, MA) generated single and double amino acid substitutions in the WF10 HA gene segment following manufacturer conditions. Plasmid sequences were confirmed by Sanger sequencing.

## Abbreviations

av: avian
dk: duck
ck: chicken
gf: guinea fowl
ty: turkey
ph: pheasant
qa: quail
ma: mallard
rt: ruddy turnstone
sh: shorebird
AR: Arkansas
DE: Delaware
NJ: New Jersey
WI: Wisconsin
Bei: Beijing
Guan: Guangdong
HK: Hong Kong
Hun: Hunan
Sic: Sichuan
Shi: Shijiazhuang
Afg: Afghanistan
Ban: Bangladesh
ENG: England
Fin: Finland
Ger: Germany
Isr: Israel
Lib: Libya
NL: Netherlands
Pak: Pakistan
RoI: Republic of Ireland
Saudi A: Saudia Arabia
S Korea: South Korea
Tun: Tunisia
UAE: United Arab Emirates
USA: United States of America

## ACKNOWLEDGMENTS

We thank the personnel from the animal resources and administrative staff at the Poultry Diagnostic and Research Center, University of Georgia, and at the Animal Plant Health Agency (APHA, UK) for technical support. Thanks to Stephen, Natalie, Susan, and James at APHA for their professional assistance. We thank Stivalis Cardenas Garcia for valuable discussions. This study was supported by a subcontract from the Center for Research on Influenza Pathogenesis (CRIP) to D.R.P. under contract HHSN272201400008C, Centers for Influenza Research and Surveillance (CEIRS) and 75N93021C00014 Centers for Influenza Research and Response (CEIRR) from the National Institute of Allergy and Infectious Diseases (NIAID). Additional funds were obtained by D.R.P under GRANT12901999, Proposal 2019-05890, Accession Number 1022658 from the National Institute of Food and Agriculture (NIFA), U.S. Department of Agriculture. D.R.P. receives additional support from the Georgia Research Alliance and the Caswell S. Eidson endowment funds from The University of Georgia. SC received a short-term training award from the NIAID CEIRS Training Program. This study was partly supported by resources and technical expertise from the Georgia Advanced Computing Resource Center, a partnership between the University of Georgia’s Office of the Vice President for Research and the Office of the Vice President for Information Technology.

## Author contributions

SC, CJC, LCG, LMF, and AO performed reverse genetics of recombinant influenza viruses, produced sera against H9 antigens in quail, and processed the samples. SC and CJC performed phylogenetic analyses. ES assisted with the initial phylogenetic analysis. DB performed initial phylogenetic analysis for the selection of consensus H9 HA antigens. IB provided H9N2 virus strains. GG sequenced viruses by NGS. DSR assisted in the interpretation of antigenic cartography. DB and MR assisted with the initial assessment of H9 HA antigens and edited the manuscript. NL assisted with antigenic cartography analyses and editing the manuscript. DRP was responsible for the overall study design, including the design of synthetic chimeric HA constructs. SC, CJC, DSR, and DRP participated in the data analysis, antigenic cartography, antigenic analysis and wrote and edited the manuscript. All authors have seen and approved the manuscript prior to submission.

